# EXTRACTION, ISOLATION, PURIFICATION AND OPTIMIZATION OF AMYLASE AND PROTEASE ENZYMES ISOLATED FROM *Bacillus* species

**DOI:** 10.1101/2025.09.10.675465

**Authors:** Aastha Acharya, Samiran Subedi

## Abstract

**Background:** Amylase and protease are two of the most demanded and widely used commercial enzymes. Amylases have many industrial uses: starch conversion (food/bakery industry), sizing agent (textile industry and paper industry), degradation of starchy residue from clothes (detergent industry), conversion of starch to fermentable sugar (fuel industry) etc. Their main source is of microbial origin. About two-third of the industrial enzymes (amylase, protease, cellulose, penicillinase, chitinase, nucleases, esterase, lipase etc.) are produced by *Bacillus* spp. Among bacteria, *Bacillus* species are specific producers of extracellular enzymes.

**Objective:** The present work comprised the identification of amylase and protease producing *Bacillus* spp and exposure of the producers to various parameters for the maximum yield of the enzyme.

**Methods:** To isolate and identify the amylase and protease producing strain, soil samples were collected from different vegetation from the altitude at 4367.35 feet above sea level. The isolates were screened and various biochemical tests and morphological observations were done to identify the isolates. The enzymes were produced by the submerged state fermentation (SmF) from the isolates and purified by dialysis. Effects of temperature, pH, and different carbon and nitrogen sources of the medium using SmF were optimized.

**Results:** Among 95 isolates, 36 were identified. Among the identified isolates, *Bacillus subtilis* and *Bacillus thuringiensis* were optimized for the amylase and protease production respectively. The maximum amylase production was found at 42⁰C temperature, in fructose as a carbon sugar, peptone as a nitrogen source and at pH 7. Similarly, the maximum protease production was found at 42⁰C temperature, in sucrose as a carbon sugar, ammonium sulphate as a nitrogen source. The producers inhabited the soil of leguminous plant.

**Conclusion:** In the present study, a natural polymer gelatin is used with the nutrient agar medium to help in cell immobilization for maximum production of alkaline protease by strains of Bacillus. More sophisticated process of purification might yield more enzyme compared to the dialysis in our process that yield 42U/ml/min and 52U/ml/min respectively.

## INTRODUCTION

Soil has been defined as the region on the earth’s crust where geology and biology meet [1]. The vast difference in the composition of soil together with the difference in the physical characteristics and agricultural practices by which they are cultivated result in the microbial population both in total numbers and kind [2]. The bacterial population of the soil exceeds the population of all other groups of microorganisms in both number and variety [3].

The metabolic abilities of bacteria play a critical role in geochemical nutrient cycling and producing a wide range of products of industrial significance. *Bacillus* is the most abundant genus in the rhizosphere [4]. The genus *Bacillus* is frequently known as aerobic spore bearers. They are ubiquitous and are present in soil, dust, air, and water. Most of the 70 or so species of *Bacillus* are found in soil and water and are usually encountered in the medical laboratory as airborne contaminants [5].The genus includes thermophilic, psychrophilic, acidophilic, alkaliphilic, halotolerant and halophilic representatives [6] which are capable of growing at temperatures, pH values, and salt concentrations at which few other organisms could survive [7]. Bacilli play major roles in the mineralization of plant-derived materials, humus, pesticides and hydrocarbons in soil [8]. Members of *Bacillus* and related genera are used for the synthesis of a very wide range of important medical, agricultural, pharmaceutical and other industrial products. Each single strain of organism produces a large number of enzymes which include the function such as hydrolyzing, oxidizing or reducing, and metabolic in nature. Amylase, lipase, protease, and cellulase constitute a very important part of microbial enzymes that are used in food, pharmaceutical, textile, paper, leather, and other industries. Most members of the genus *Bacillus* are able to produce low molecular weight antibiotics, which possess different biological activities, including antimicrobial, antiviral, and antitumor activities [9].

Enzymes may be membrane bound in young cells and released as exoenzymes as the culture enters stationary phase [10] or solubilized by relatively mild procedures including washing the cells with water or concentrated salt solution [11]. The amylases and proteases are probably the most ubiquitous.

Amylase is an enzyme that catalyses the hydrolysis of starch by breaking the bonds between sugar molecules. It can be found in animals, plants, and bacteria. Amylase can be classified into three types: α-amylase (EC 3.2.1.1), β-amylase (EC 3.2.1.2) and γ-amylase (EC 3.2.1.3) [12]. Amylases are produced by a variety of living organism, ranging from prokaryotes (bacteria) to eukaryotes (plants and animals). Many microorganisms have a potential to produce amylases including *Bacillus subitilis, Bacillus megaterium, Bacillus cereus, Lactobacillus* spp, *Escherichia coli*, *Pseudomonas* spp, *Streptomyces* spp, etc [13].

Protease (also called peptidase or proteinase) is an enzyme that performs proteolysis (protein catabolism by the hydrolysis of the peptide bonds). It can be found in the animal, plant, fungi, bacteria, archae and actinomycetes. Proteases can be classified in many types: serine proteases, cysteine proteases, aspartic proteases, threonine proteases, glutamic proteases, metalloproteases and asparagine peptide lyases [14]. Group A serine protease is typified by the enzymes from *B. licheniformis* and *B. pumilus*; and group B serine proteases are related to *B. subtilis* in novo, the enzyme from *B. amyloliquefaciens*. *B. megaterium*, *B. polymyxa*, *B. thermoproteolyticus* and *B. thuringiensis* secrete only the metal protease [41]. Several *Bacillus* species involved in protease production are *B. cereus*, *B. sterothermophilus*, *B. mojavensis*, *B. megaterium* and *B. subtilis* [17].

Different strategies for purification of enzymes have been investigated, exploiting specific characteristics of the target biomolecule [18]. Enzyme purification can be done by using the following techniques: Ammonium Sulphate, Precipitation, Dialysis, Ultra filtration and Lyophilization [17]. The optimization of fermentation conditions, particularly physical and chemical parameters, are important in the development of fermentation processes due to their impact on the economy and practicability of the process [19].The nature and amount of carbon sources in culture media are important factor for the production of extracellular enzymes [14]. Temperature is a vital environmental factor which controls the growth and production of metabolites by microorganisms and this is usually varied from one organism to another [20]. Enzyme production was found to gradually increase with increasing temperature and maximum enzyme production was observed at 42⁰C. Among physical parameters, pH of the growth medium plays an important role in enzyme secretion [21]. The pH range observed during the growth of microbes also affects product stability in the medium [22].

This study is undertaken to isolate Bacillus from the rhizospheric region of various vegetations, and assess their ability to produce protease and amylase enzymes. The purpose of this study is to investigate the effectiveness of production of enzyme (protease and amylase) by incubating test *Bacillus* spp in media differing in one of the aspects: Carbon source, Nitrogen source, pH, temperature. The set of conditions which support higher degree of enzyme production can be used for more economical and higher yield of enzyme.

## MATERIALS and METHODS

### Study Area

The study area for sample collection was from Kathmandu valley which was located at an altitude of 4367.35 feet above sea level. Soil was collected using random sampling method from different places of Kathmandu.

### Study Method

Random sampling method was applied for the collection of the samples. Soil samples were collected from various wet and dry land.

### Sample Size

A total of 25 samples were collected from different vegetations of Balaju Nursery House which is at an altitude of 4367.35 feet above sea level.

### Study Duration

The project was carried out in microbiology laboratory of St. Xavier’s College, Maitighar, Kathmandu, Nepal from 15th October, 2016 to 10th June, 2017.

### Sample Collection

Soil samples were collected in an aluminum foil from a depth of 5 cm.

### Isolation of Bacteria from Soil

One gram of soil sample was taken and serially diluted up to 10^−6^ in sterile distilled water and spread plate technique was performed for the isolation of bacteria. For this 0.1 ml of sample was dispensed in petri plate containing nutrient agar and HiCrome *Bacillus* agar and was spread using sterile glass rod. Samples from dilution 10^−6^ were subjected to spread plate into respective plates. The plates were then incubated at temperature 28°C for 24 hours [23].

### Primary Screening of Protease Producer

Different isolated colonies from the nutrient agar plates were chosen and sub cultured in 1% Gelatin agar plate and some bacterial isolates were sub cultured in nutrient broth as stock culture and were incubated at temperature 28°C for 24 hours. After incubation period, the gelatin agar plates were then flooded with 15% HgCl_2_ –HCl solution. HgCl_2_ was then dispensed with caution and zone of hydrolysis was observed [23].

### Primary screening of Amylase Producer

Different isolated colonies from the nutrient agar plates were chosen and sub cultured in 1% Starch agar plate and some bacterial isolates were sub cultured in nutrient broth as stock culture and was incubated at temperature 28°C for 24 hours. After incubation, the starch agar plate was flooded with iodine solution. Iodine was then dispensed with caution and zone of hydrolysis was observed [23].

### Characterization of Protease and Amylase Producer

Gelatin and starch hydrolyzing bacterial isolates were then characterized by using standard microbiological technique which included colony morphology, Gram’s stain and biochemical reactions and pure colonies were examined for *Bacillus* with standard description of Bergey’s Manual of Determinative Bacteriology.

### Production of Extracellular Enzyme (Submerged Fermentation)

The gelatin and starch hydrolyzing isolates were inoculated in conical flask containing 50 ml of 1% gelatin broth and 1% starch broth respectively. The flasks were then kept in shake flask rotary incubator. Time, temperature and speed (in rpm) were set at 24 hour, 28°C and 120 rpm respectively.

### Extraction of Extracellular Protease

The broth cultures were transferred in tubes and labeled. It was centrifuged at 2000 rpm for 20 minutes and the supernatant was transferred to another test tube and pellets were discarded. The tube containing supernatant was mixed with double volume of acetone and was left overnight and again centrifuged at 3000 rpm for 20 minutes.

Supernatant was transferred to other test tube(s) and pellet was dissolved in minimum volume of phosphate buffer. These enzymes were labeled as crude enzyme [24].

### Extraction of Extracellular Amylase

The broth cultures were transferred in tubes and labeled. It was centrifuged at 2000 rpm for 20 minutes and the obtained supernatant was transferred to another test tube and pellets were discarded. The tube containing supernatant was mixed with double volume of acetone and was left overnight and again centrifuged at 3000 rpm for 20 minutes.

Supernatant was transferred to other test tube(s) and pellet was dissolved in minimum volume of phosphate buffer. These enzymes were labeled as crude enzyme [24].

### Purification of Crude Enzyme

The crude enzyme obtained was purified by using dialysis bag. The crude enzyme was kept in the dialysis bag and was introduced in a beaker with distilled water and purification was done using magnetic stirrer for 40 minutes.

### Determination of Enzyme Activity

#### For Protease

Proteolytic activity was measured by using casein as a substrate. 0.5% casein (2ml) was added to the crude enzyme obtained from acetone fractionation and incubated for 30 minutes at 50°C.

The mixture was centrifuged at 2000 rpm at 4°C for 10 min. The clear supernatant was mixed with 0.5 ml of 1N Folin Ciocalteau’s reagent and the absorbance was measured at 680 nm.

#### For Amylase

Amylase activity in the crude enzyme was determined by estimation of the reducing sugar liberated by the action of amylase on soluble starch. Reducing sugars were estimated by 3,5 – Dinitrosalicylic acid (DNS) method [25].

### Enzyme optimization

For enzyme optimization, multiple sets of conical flask-each containing 50 ml broth media differing uniquely in carbon, nitrogen, pH and temperature and time of incubation were inoculated with test organism. The enzyme so produced were assayed using corresponding methods as described above.

### Quality control

Laboratory equipments like incubator, hot air oven, autoclave, refrigerator, etc. were regularly monitored for their performance and immediately corrected if any deviation occurred. The temperature of the incubator and refrigerator were mentioned every day.

For consistency in results, any reagents and media were prepared fresh and stored under appropriate conditions. Any media or reagents to be used were prepared in sterile (if not aseptic) condition.

### Data analysis

Data were statistically analyzed using various graphs and tables.

### Validity and reliability

For the valid test results standard protocol was followed and pre-testing of the tests was performed. Similarly, reliability of the study depended on expert’s opinion, guidance of supervisor’s and literature review.

### Limitation of the study

The isolated *Bacillus* species may not include all the protease and amylase producers as the soil sample were collected from limited areas. There lies a difficulty in identification of species as molecular techniques are not available.

## RESULT

All together 25 soil samples were taken from the rhizospheric region of different vegetations from the Balaju Nursery which is at an altitude of 4367.35 feet above sea level. Of the 95 isolates obtained, 36 species were identified. Of the 36 species isolated *B. polymyxa* was observed to be the most prominent one with overall of 20% population followed by *B. circulans* with 14%, *B. megaterium* 11%, *B. coagulans* and *B. licheniformis* with 9%. Similarly 60% bacterial isolates were identified as protease producer and 73% as amylase producer among the isolates. From the identified species, *B. subtilis* and *B. thuringiensis* were optimized for the amylase and protease production under different parameters.

### Identification of the isolates

Based on the color of colonies observed on HiCrome *Bacillus* agar [28] and the biochemical tests based on the Bergey’s Manual of Determinative Microbiology [26], the isolated colonies were identified. Since, the identification of *B. thuringiensis* was not mention in the manual, for its detection, biochemical tests were carried out based on the procedure described [27]. Similarly, for the identifaction and detection of *Bacillus thuringensis* among the isolates, the crystal protein staining was also performed as outlined by Sharif and Alaeddinoĝlu [29]. The results are presented in table 1.

**Table 1:**
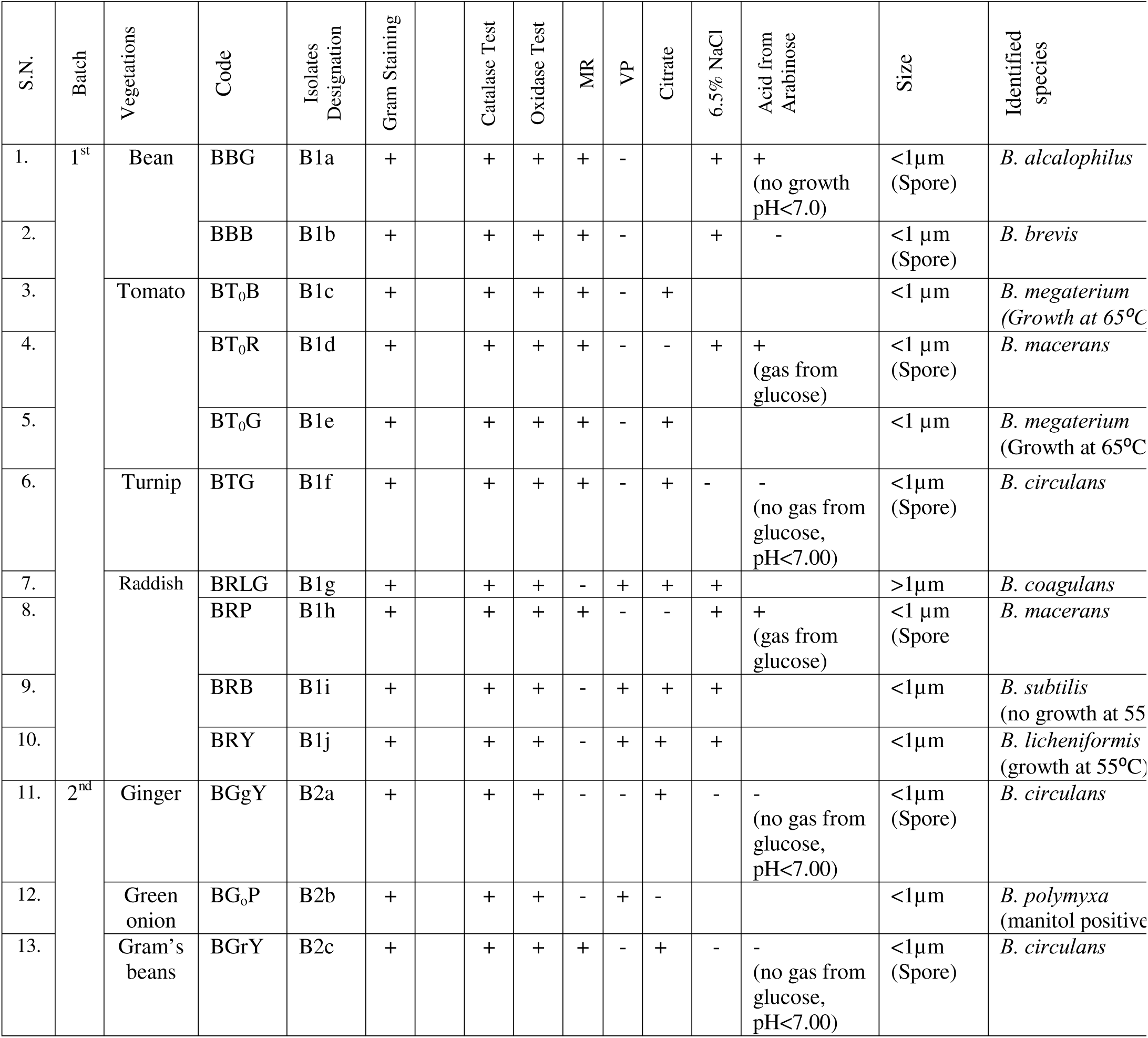
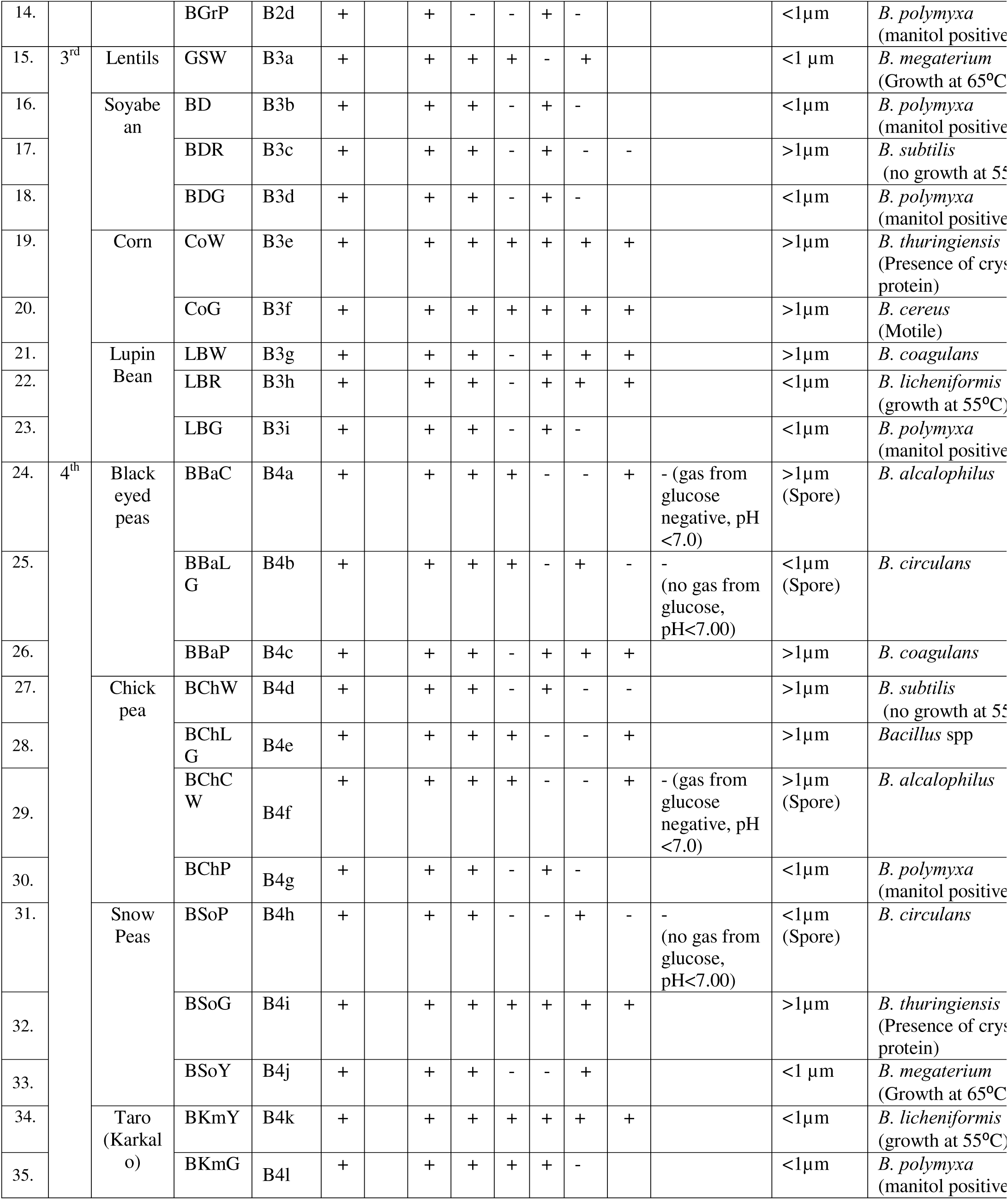
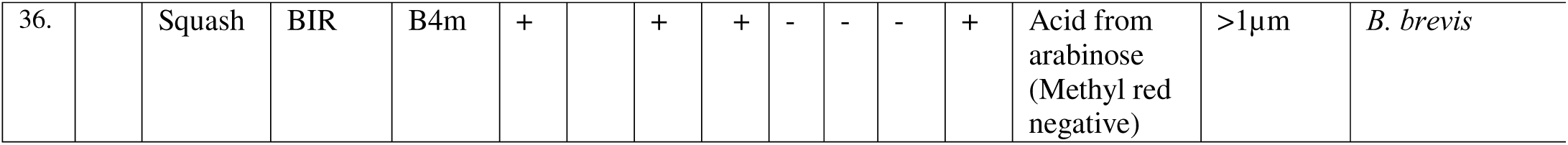
Identification of species. Bacterial colonies observed on HiCrome *Bacillus* Agar were further identified by the biochemical test such as methyl red (MR), Vogues-Prausker (VP), Catalse, Oxidase, Citrate, Acid from Arabinose test along with micrometry and their ability to grow om 6.5% NaCl, mannitol production and motility test as described on Bergey’s Manual of Determinative Microbiology.

### Distribution of isolated species

A total of 36 species were identified and are presented in the figure below Figure 1. Identification of selected *Bacillus* strain was on the basis of standard morphological and biochemical test according to the method described in Bergey’s Manual of Derterminative Bacteriology Highest amylase production was seen in Badam and protease production was seen in green vegetable (Figure 1).

**Figure 1:**
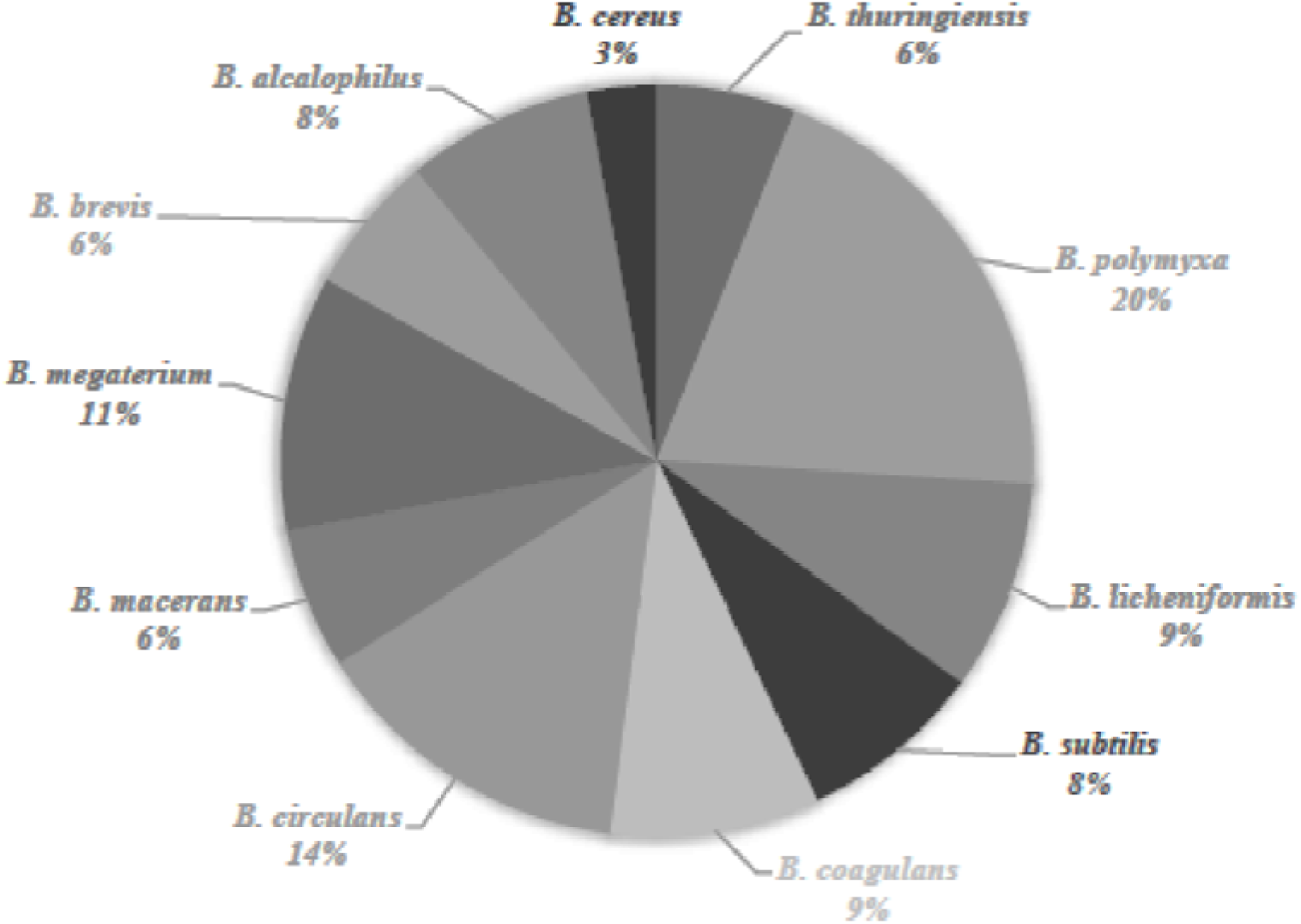
Population of the each species among the total 36 identified species

### Assay of protease produced by *Bacillus thuringiensis* under different parameters

For further studies on production of protease, shake-flask culture was done using *Bacillus thuringiensis*. This organism was selected for protease production as it yield the maximum protease as compared to other *Bacillus* spp. The production was carried out in the basal medium where the sugars (glucose, maltose, lactose and sucrose), nitrogen source (yeast extract, peptone, urea and ammonium sulphate) were used at different temperatures (20⁰C, 28⁰C, 32⁰C, 37⁰C and 42⁰C) and pH 7 where Casein (0.5%) was used as a substrate.

When glucose was used in conjugation with different nitrogen sources, different readings were observed. The maximum absorbance was observed when glucose was used with peptone (absorbance of 1.15), followed by urea (1.09) and least in case of ammonium sulphate with the reading at 0.82 (Figure 2).

**Figure 2:**
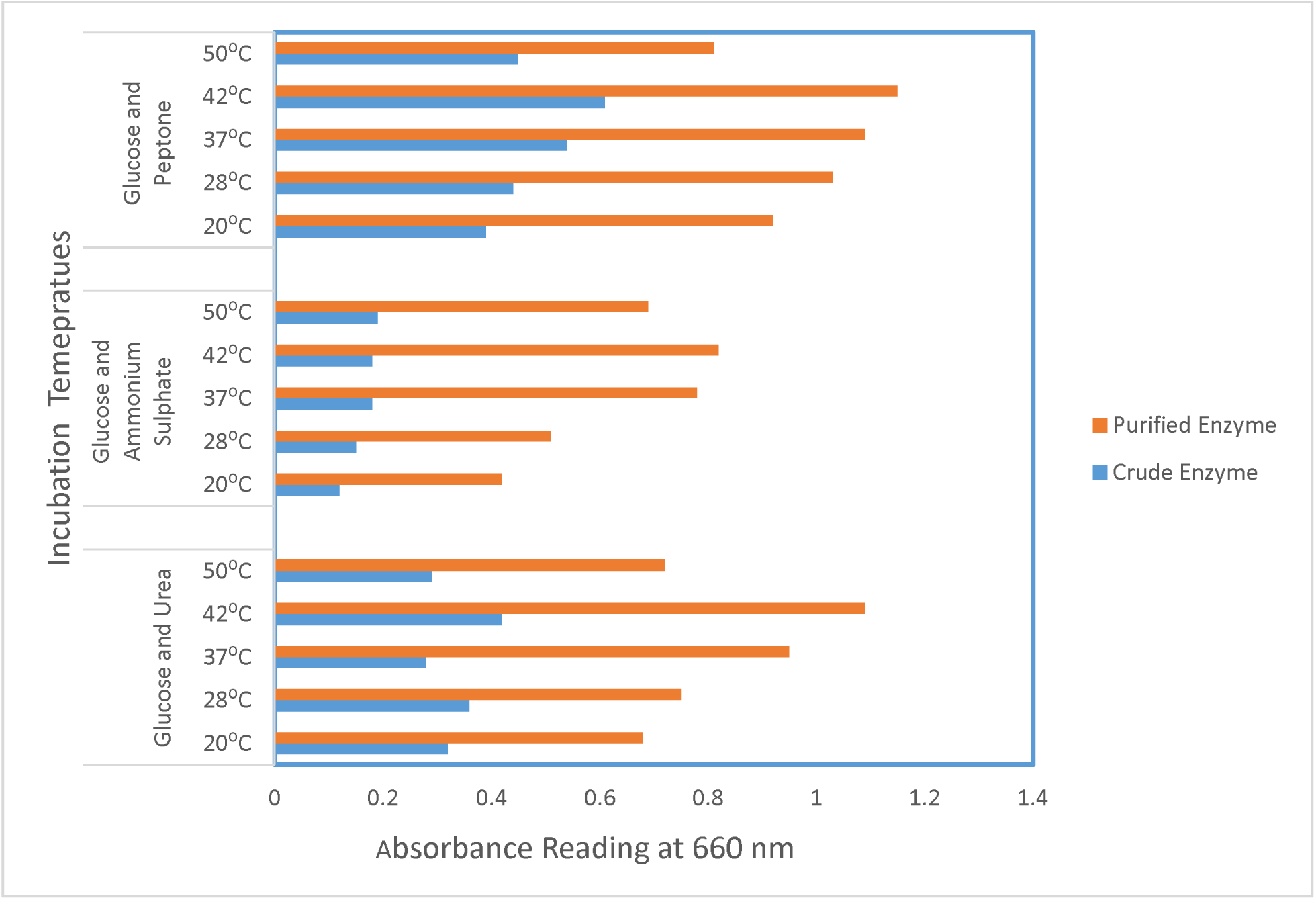
Protease activity of *Bacillus thuringiensis* upon incubation with Glucose and different Nitrogen Sources (Peptone, Ammonium Sulphate and Urea) under various incubation temperatures.

Similarly, sucrose was used as a carbon source along with peptone, ammonium sulphate and urea as a nitrogen source respectively. The maximum result was obtained when sucrose was used in amalgamation with ammonium sulphate that gave 1.72 followed by peptone, 1.37 and urea with 0.58 µ/ml respectively (Figure 3)

**Figure 3:**
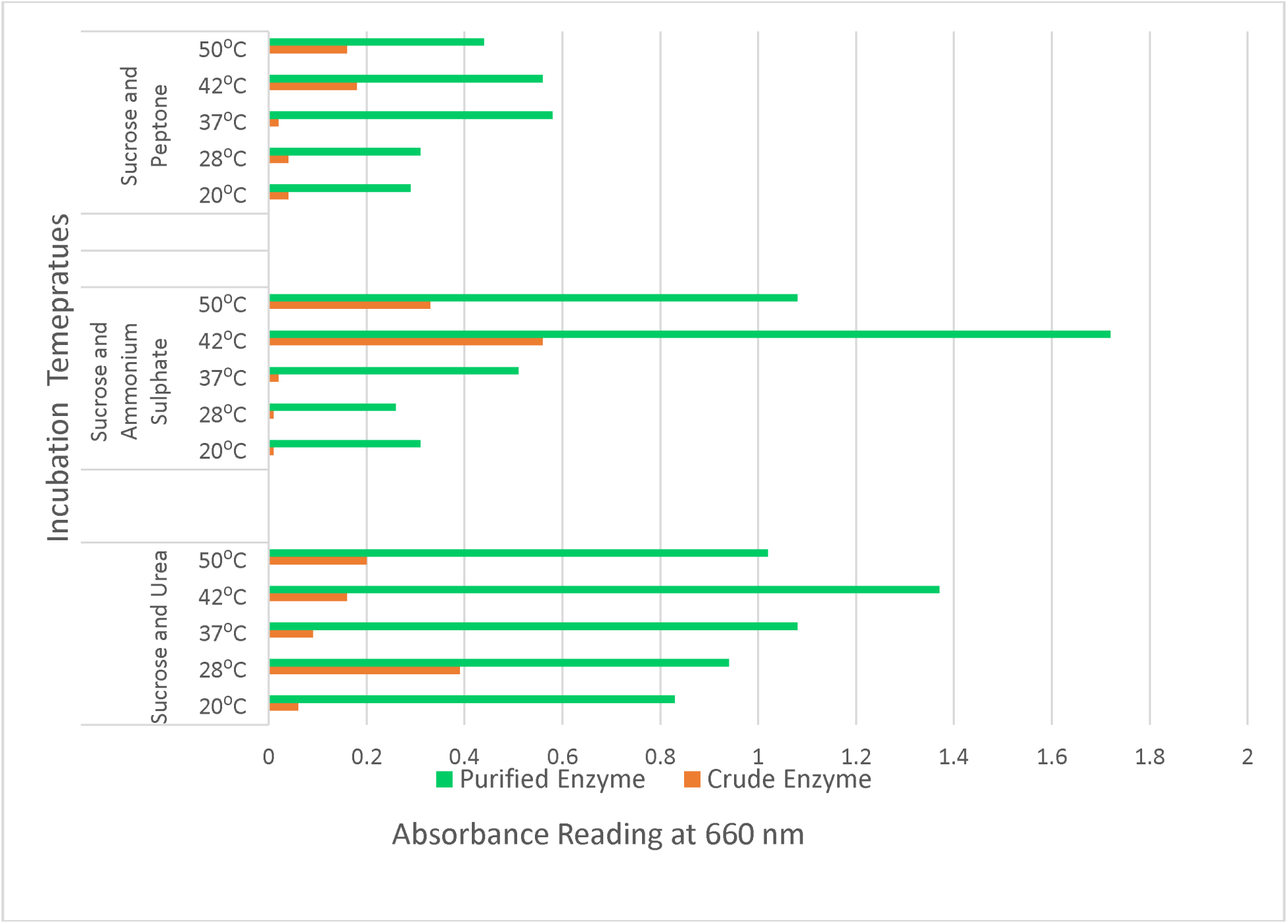
Protease activity of Bacillus thuringiensis upon incubation with Sucrose and different Nitrogen Sources (Peptone, Ammonium Sulphate and Urea) under various incubation temperatures.

Likewise when maltose was used with the different forms of nitrogen source, the maximum absorbance was observed in case of peptone with 1.76 µ/ml at 37⁰C followed by ammonium sulphate with 0.97 µ/ml at 42⁰C and urea with 0.83 µ/ml at 42⁰C (Figure 4)

**Figure 4:**
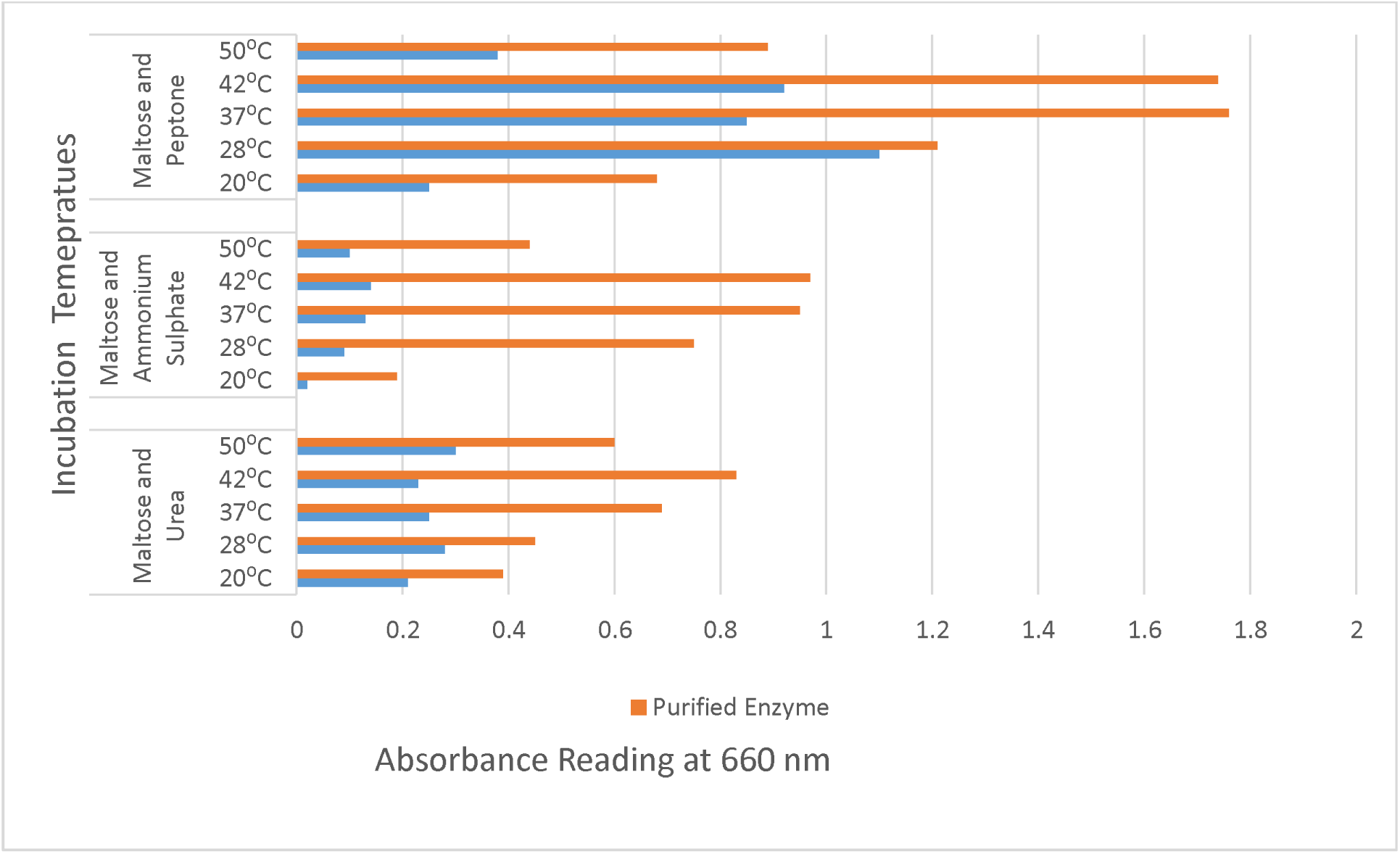
Protease activity of Bacillus thuringiensis upon incubation with Maltose and different Nitrogen Sources (Peptone, Ammonium Sulphate and Urea) under various incubation temperatures.

Correspondingly, when lactose was used as carbon source with different nitrogen sources such as peptone, ammonium sulphate and urea respectively, the maximum reading was observed in case of urea which was indicated by maximum absorbance of 1.93 followed by peptone with absorbance 1.25 and ammonium sulphate with absorbance of 0.82. (Figure 5)

**Figure 5:**
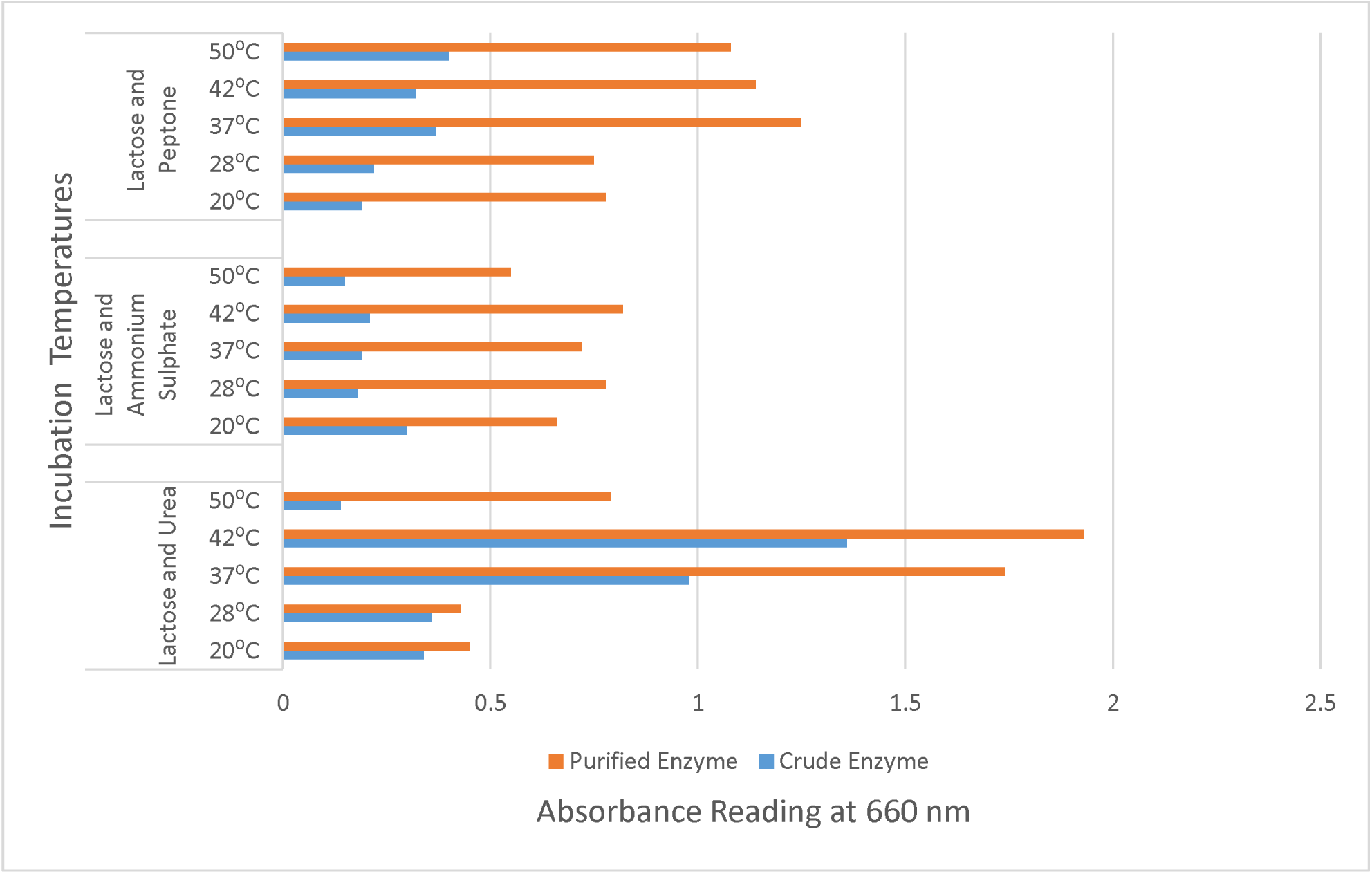
Protease activity of Bacillus thuringiensis upon incubation with Lactose and different Nitrogen Sources (Peptone, Ammonium Sulphate and Urea) under various incubation temperatures.

The optimization of simpler form of nitrogen source i.e yeast extract was observed in conjugation with different forms of carbon source such as glucose, sucrose, lactose and maltose respectively. The optical density in each of the case were observed and maximum result was found in case of lactose and maltose in conjugation with yeast extract at 37⁰C with the highest absorbance of 1.3 (Figure 6)

**Figure 6:**
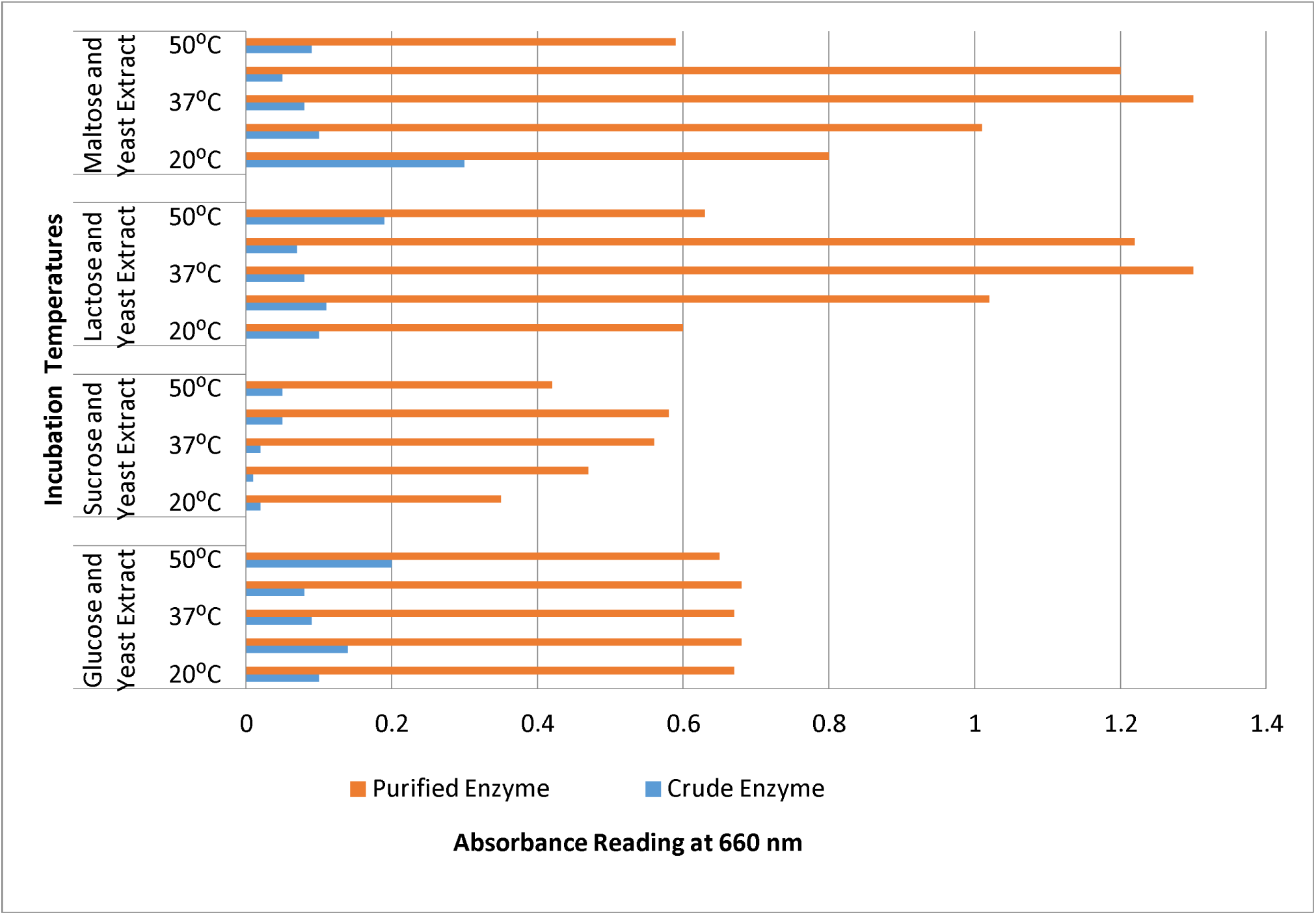
Protease activity of Bacillus thuringiensis upon incubation with Yeast and different Carbon Sources (Glucose, Sucrose, Lactose and Maltose) under various incubation temperatures.

### Assay of amylase produced by *Bacillus subtilis* under different parameters

The genus *Bacillus* produces a large variety of extracellular enzymes, of which amylases are of particularly considerable industrial importance [42]. Since *Bacillus subtilis* yield the maximum amylase production among other organism, it was associated with various parameters in shake-flask culture using starch (0.5%) as a substrate along with different forms of sugars (glucose, fructose, lactose and sucrose), nitrogen source (peptone, urea and ammonium sulphate) at different temperatures (27⁰C, 32⁰C,37⁰C and 42⁰C) and pH 7.

For the optimization of amylase, the organism was inoculated in media with variety of nitrogen source (peptone, urea and ammonium sulphate) coupled with various carbon source (glucose, fructose, sucrose and lactose) and incubated under various temperatures. Peptone was considered suitable nitrogen source as compared to urea and ammonium sulphate according to the optical density readings at 540 nm (Figure 7). Similar readings were observed in case of fructose where maximum reading was observed when peptone was used along with fructose compare to other two nitrogen sources (Figure 8).

**Figure 7:**
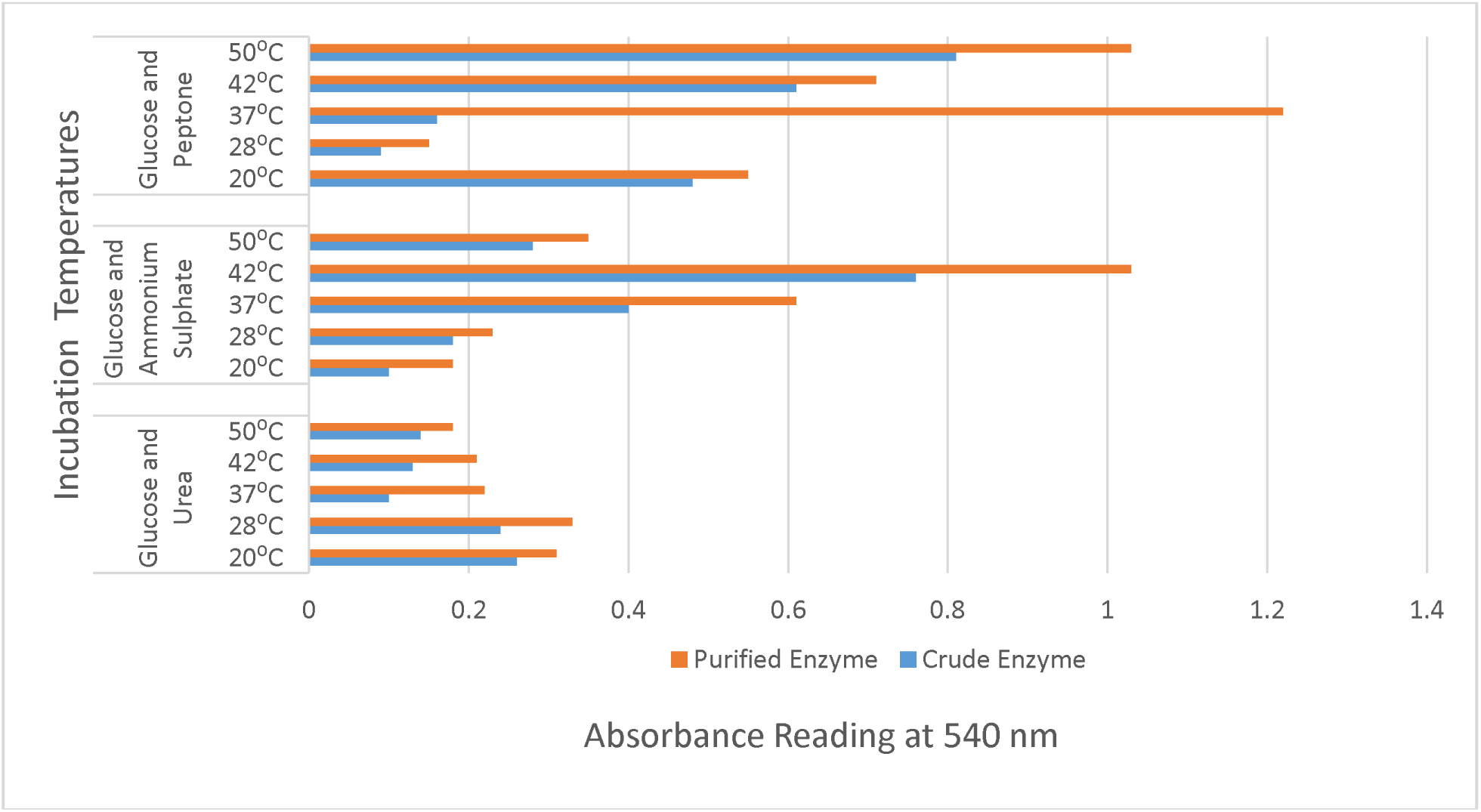
Amylase activity of Bacillus subtilis upon incubation with Glucose and different Nitrogen Sources (Peptone, Ammonium Sulphate and Urea) under various incubation temperatures.

**Figure 8:**
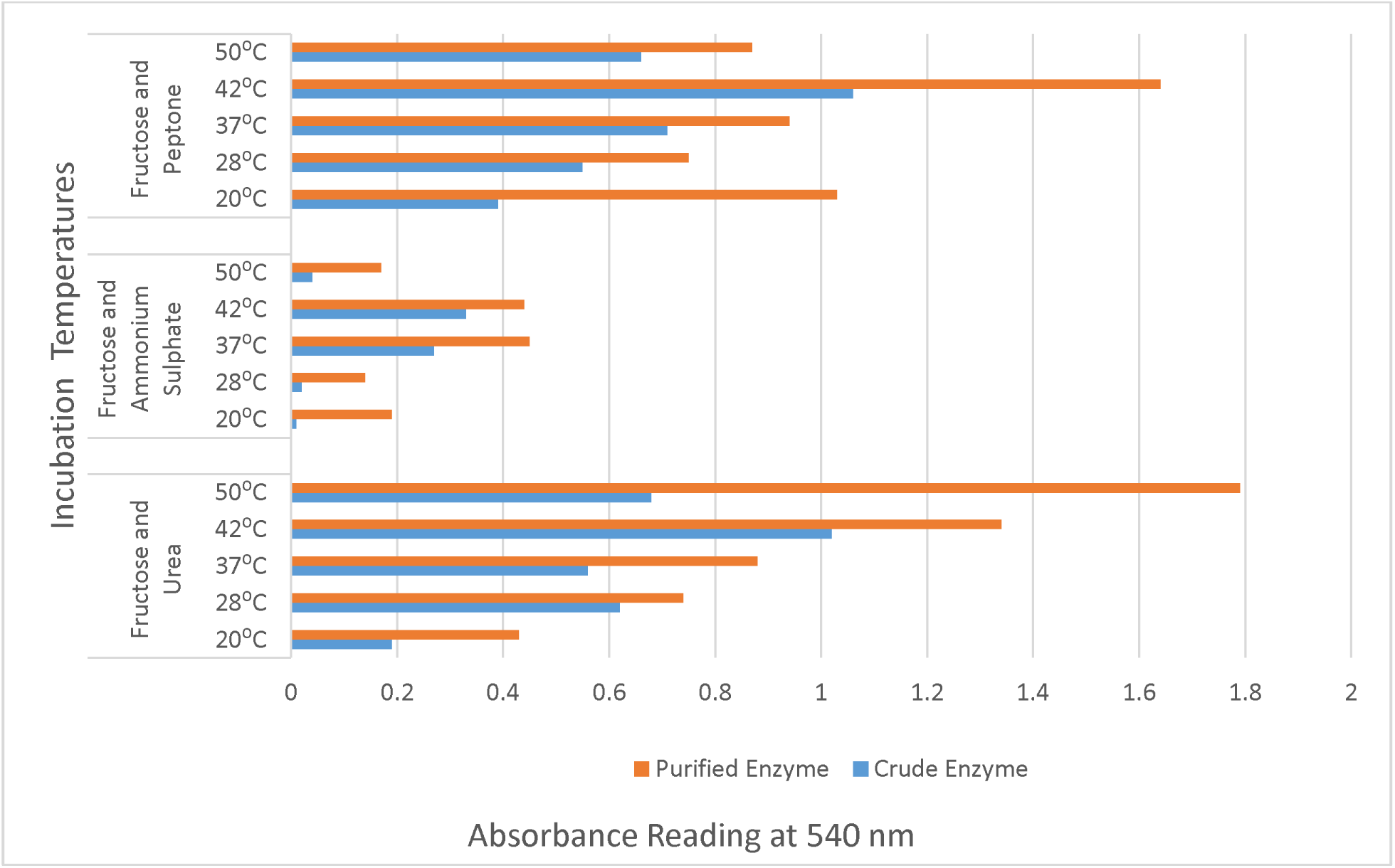
Amylase activity of Bacillus subtilis upon incubation with Fructose and different Nitrogen Sources (Peptone, Ammonium Sulphate and Urea) under various incubation temperatures.

Comparably, the optimization of the different nitrogen source was observed with lactose and sucrose respectively. Lactose corporate media with the different nitrogen sources gave similar reading at all temperatures (Figure 9). Whereas, sucrose and urea gave the maximum result at 42⁰C. (Figure 10).

**Figure 9:**
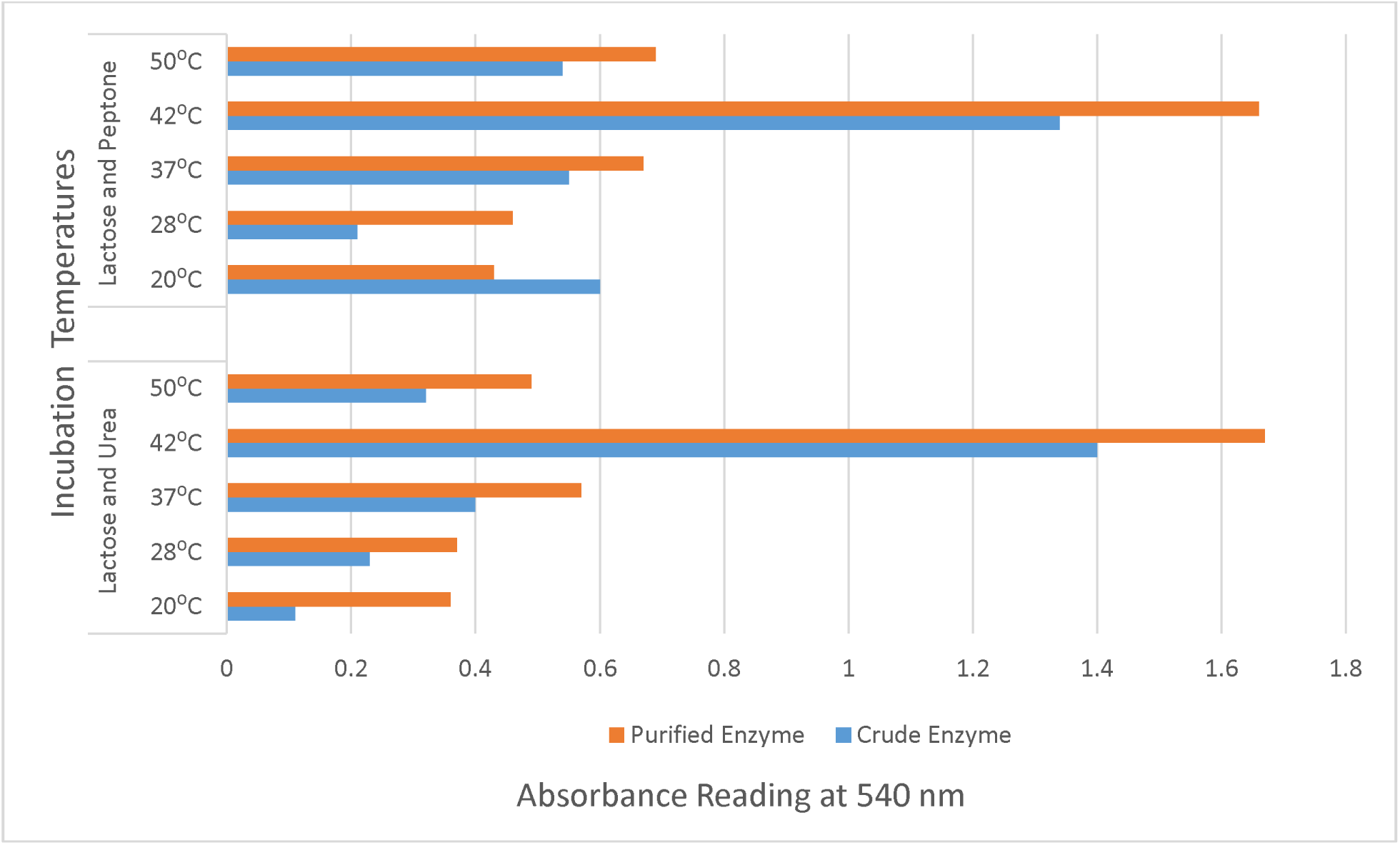
Amylase activity of Bacillus subtilis upon incubation with Lactose and different Nitrogen Sources (Peptone and Urea) under various incubation temperatures.

**Figure 10:**
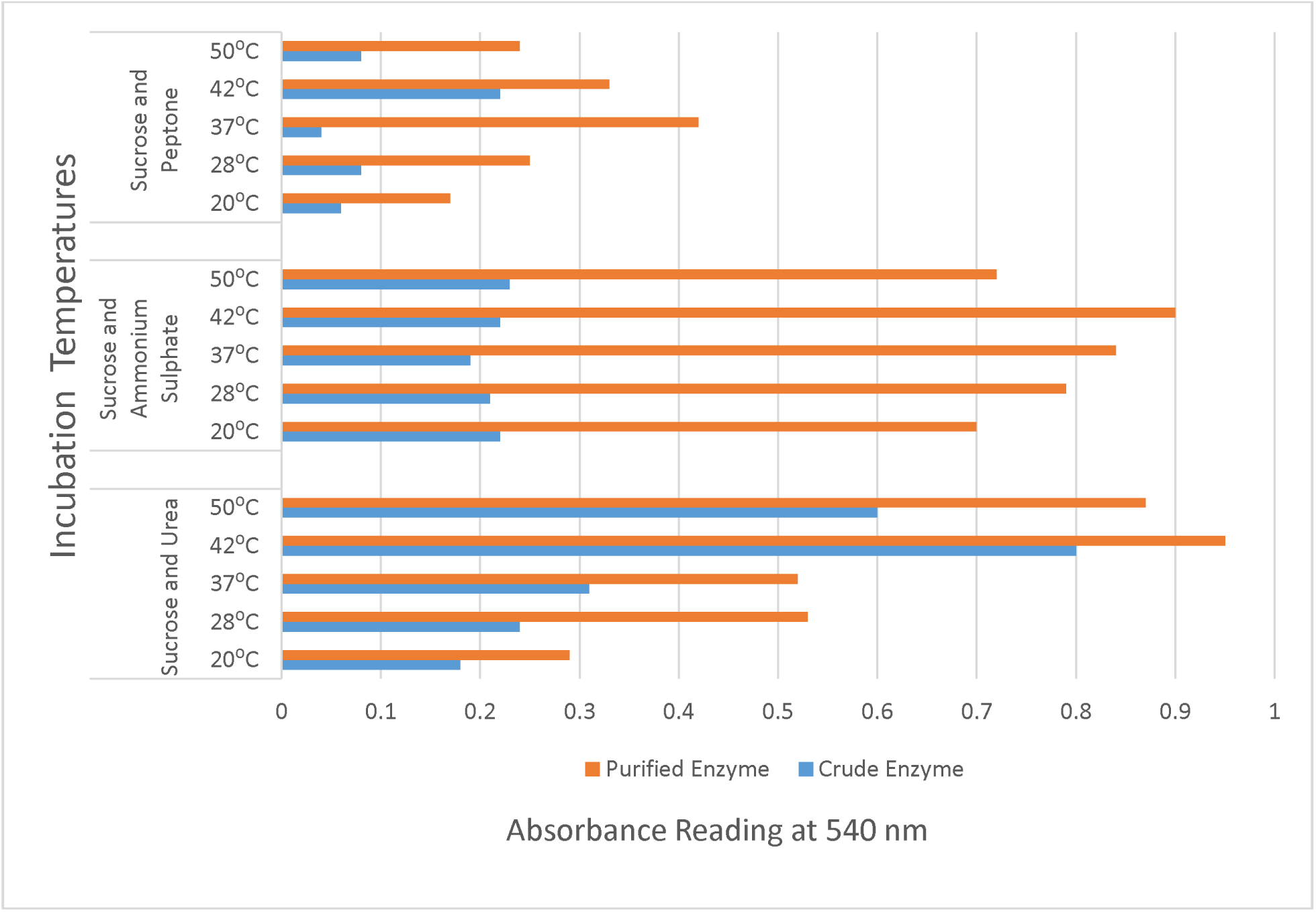
Amylase activity of Bacillus subtilis upon incubation with Sucrose and different Nitrogen Sources (Peptone, Ammonium Sulphate and Urea) under various incubation temperatures.

Among various pH ranges (3.6, 7.0 and 9.0), pH 7 was considered to be the best both in amylase and protease production in presence of glucose and sucrose as carbon source along with peptone as a nitrogen source at 37⁰C which are shown in figures 11 and 12 respectively.

**Figure 11:**
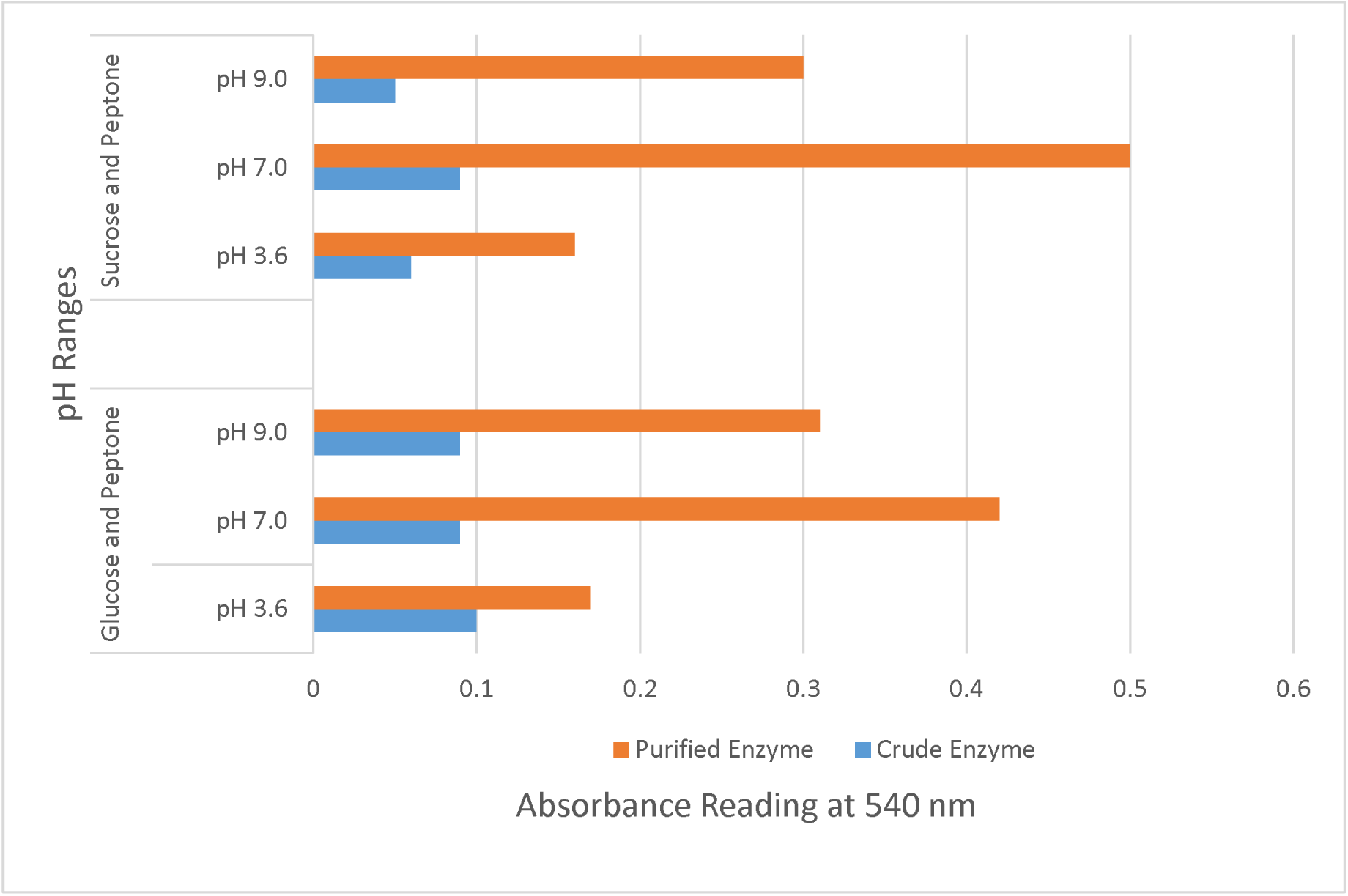
Amylase activity of Bacillus subtilis upon incubation with Carbon source (Glucose and Sucrose) and Peptone under various pH.

**Figure 12:**
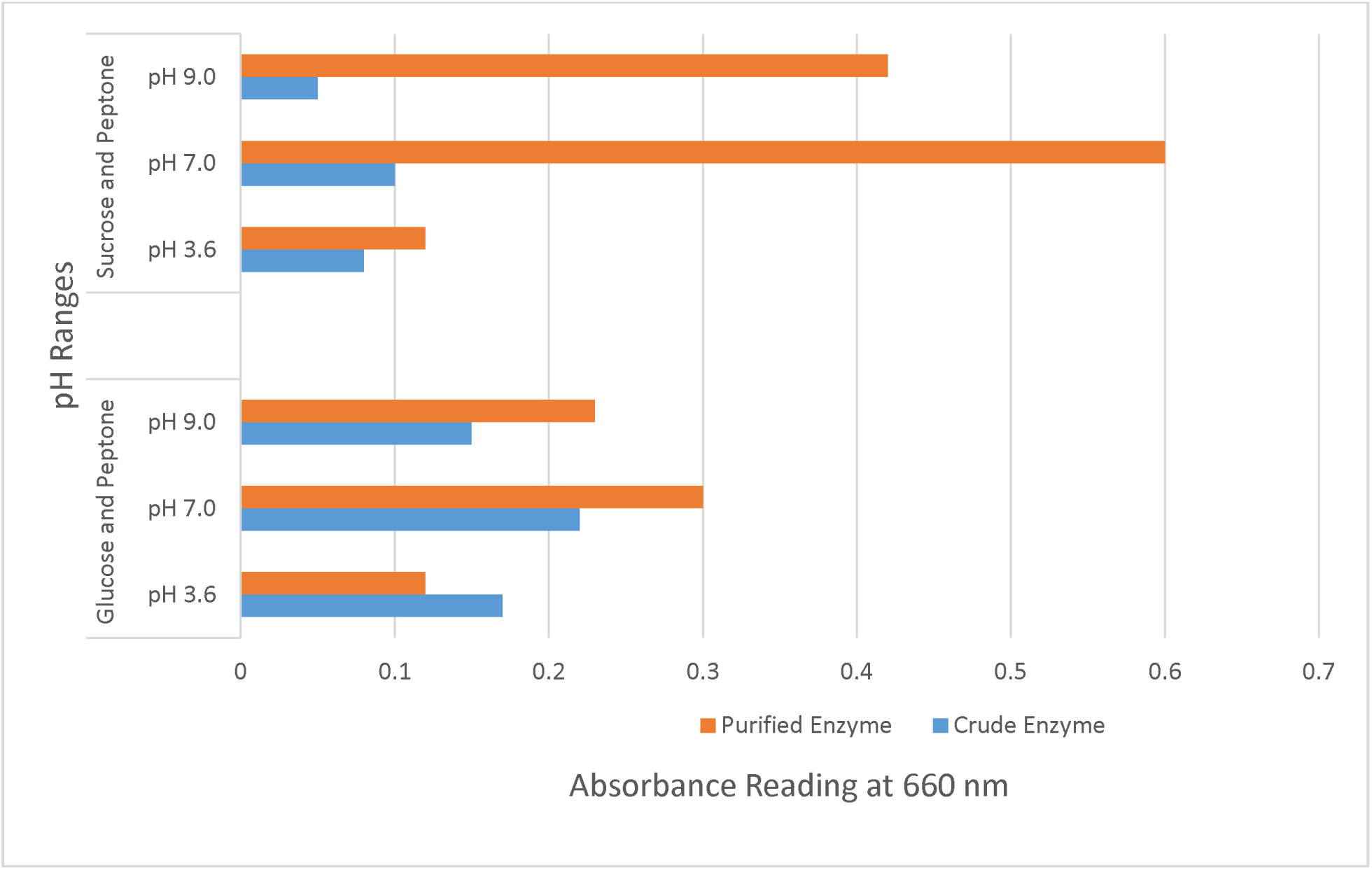
Protease activity of Bacillus thuringiensis upon incubation with Carbon source (Glucose and Sucrose) and Peptone under various pH.

## DISCUSSION

This study shows the potential of *Bacillus* species to produce highly commercially important enzyme capable of starch and protein degradation.

Enzyme producing microorganisms are abundantly present in the rhizospheric region especially of the leguminous plant. Various enzymes are present in the root exudates. *Bacillus* is the most abundant genus in the rhizosphere [4]. Bacilli such as *B. subtilis*, *B. cereus*, and *B. mycoides* are known for their roles as beneficial rhizobacteria that promote plant growth [30].The colonies on HiCrome *Bacillus* Agar were characterized on the basis of color. A similar approach has been done by [28].

A total of 95 isolates were obtained from the samples of which 60% bacterial isolates were identified as protease producer and 73% as amylase producer. This activity was seen by development of zone of hydrolysis by growth of organism in NA media with 1% of respective substrate. The most frequently occurring amylolytic bacteria with greatest enzyme activity was *Bacillus subtilis* followed by *B. polymyxa, B. megaterium, B. coagulans, B. licheniformis* and *B. cereus.* In a similar study performed by [11] *Bacillus subtilis* was found to be most frequently occurring amylase producer. The highest proteolytic activity was shown by *B. thuringiensis* followed by *B. subtilis*, *B. licheniformis* and *B. polymyxa*.

Enzyme production was carried out by submerged fermentation in starch and gelatin broth for the amylase and protease production respectively. Crude enzyme obtained after centrifugation was purified via dialysis. The activity of the purified amylase enzyme was detected via the Bernfeld method using DNS reagent [32]. Absorbance of color compound was measured at 540 nm in colorimeter. Similarly, for the protease, the activity of purified enzyme was detected via the Lowry’s method in 660 nm as depicted by [43]. Optimal culture conditions such as sources of nitrogen and carbon, their ratio, pH, temperature of incubation, etc. ensure maximum growth, metabolism and production of enzyme. Thus, optimization of growth condition is a prime step in fermentation technology. In this study, we observed 42°C as the optimum temperature for the enzyme production. This may be because of mesophilic nature of the organism. Our finding correlates with the study carried out by [33].

The assay of amylase activity was done using DNS method which is based on the reduction of 3,5-dinitrosalicylicaacid by reducing sugar to 3-amino-5-nitrosalicylic acid. The reaction imparts orange red color whose intensity depends on concentration of sugar and can be read colorimetrically at 520-540nm.

When glucose was used as a carbon source with respective nitrogen sources: urea, ammonium sulphate and peptone, varying results were observed. These variations in the results is given in the theory proposed by Szilard [44] which suggests that when there is an external manipulation in the enzyme system (in our case the temperature) when glucose is used as a carbon source, the maximum activity is seen at those temperature that favors the enzyme production irrespective of the nitrogen sources used.

Lactose with peptone had the absorbance of 1.66 whereas with urea, it was 1.67. This is because both the peptone and urea are simpler nitrogen that can be easily assimilated. The transportation and breakdown of lactose is regulated via the lac operon system. Due to the similar lactose degradation pathway and ease of assimilation of peptone and urea, the absorbance in both cases were found to be more or less similar.

Though fructose and glucose, both are simpler form of carbohydates, fructose yielded more effective and fruitful results. The maximum enzyme activity was observed at 42⁰C with absorbance of 1.67 when it was used along peptone. This was unlike glucose where variable results were found. This is because, fructose is comparatively fermented faster than glucose because it skips one step in glycolysis and when it acts along with simpler form of nitrogen, it tends to give more efficient result. This agreed with the findings of Suribabu [34].

Most of the starch degrading bacterial strain revealed a pH range between 6.0 and 7.0 for normal growth and enzyme activity [14]. The optimum pH was found to be 7 which showed optimum enzyme activity despite the different sugars that were used. The finding accord with the study done by Devi [15]. The nitrogen source, when it is utilized by bacteria, they break it down into simpler product and increase the pH ranges and thus increase the enzyme activity. This is evident from the facts that maximum enzyme activity was seen at pH 7 in all cases. Enzyme production was comparatively higher at pH 9 than at pH 3.6.

The assay of protease activity was done using Lowry’s method which is based on the reaction between the aromatic amino acids present in the sample with Biuret reagent, followed by a reaction with Folin-phenol reagent. The reaction imparts dark blue/ purple color to the solution whose intensity can be read colorimetric ally at 660-680 nm [16]. More the protease activity, more are the free proteins and aromatic acids and thus more is the color intensity (and the value of absorbance). Since everything is identical, including the test organism, its load, concentration of substrate and time of incubation; the scenarios where high values of absorbance is observed represents such parameters which favor higher production of protease.

The optimum temperature for enzyme production by *Bacillus thuringiensis* is 40-50 ⁰C [35]. In our study, the optimum temperature for protease production, in most cases, was found to be at 42°C when glucose, sucrose, maltose and lactose were used as carbon source. When glucose was used as carbon source, the pairing of simple organic nitrogen sources like peptone or urea resulted in higher production of protease than the pairing of inorganic or complex organic nitrogen sources like ammonium sulphate and yeast extract respectively. This is because urea and peptone are comparatively simpler and can be easily assimilated by the bacteria. This is in accordance with findings of [36] who stated that yeast extract along with the complex carbon sources is found to be the good inducer of the alkaline enzyme production. Since glucose concentration is 1% and yeast extract is 0.5%, the findings yet again correlate with findings of [36]: higher concentrations of glucose than yeast promote the enzyme production regardless of the incubation temperature. High concentrations of glucose and yeast extract supported better growth and biomass but enzyme production was suppressed at glucose concentrations above 1%.

Use of sucrose in combination with ammonium sulphate gave very high absorbance reading at 42°C. Sucrose, being disaccharide, is hydrolyzed into glucose and fructose-both of which can enter EMP pathway with even fructose requiring one less step to do so. The ease of assimilation of sugars along with high concentration of ammonium ions help facilitate glutamate dehydrogenase and synthase pathways that eventually lead to higher rates of creation of amino acids and α keto glutarate to be used in Kreb’s cycle [37, 38].

The combination of sucrose with peptone led to higher production of protease than that obtained from glucose and peptone [39]. Since sucrase activity is optimum at 37°C, higher assimilation occurs at this temperature which might have upper hand in production of protease when complex nitrogen source like yeast extract is used.

Upon pairing maltose with nitrogen sources, the overall production of protease was found to be little higher that that obtained by pairing glucose and corresponding nitrogen sources. Because one molecule of water is lost upon glycosidic bond formation, hydrolysis of maltose consequently leads to higher number of glucose units (theoretically 1glucose more per 10 glycosidic bonds). Lactose was also found to be potent carbon source for production of protease however, the rates of influx of lactose inside bacterial cell is based on the reactivity and optima conditions of β-galactosidase enzyme produces as a result of inducible lac operon system. As a result, the assimilation happens to be the major rate limiting step. The optima of β-galactosidase being at 37°C means that the rate of assimilation and ultimately production of protease enzyme will tend to be higher at 37°C. The production of maximum enzyme at 37°C maybe also be due to the mesophilic nature of *Bacillus* as stated by Yadav [40].

## CONCLUSION

The use of amylase and protease in industries has been prevalent for many decades and a number of microbial sources exist for the efficient production of this enzyme, but only a few selected strains of fungi and bacteria meet the criteria for commercial production. The search for new microorganisms that can be used for amylase production is a continuous process. More recently, many authors have presented good results in developing enzyme purification techniques, which enable applications in pharmaceutical and clinical sectors which require high purity amylases. Bacillus species can be used for large-scale production of alkaline enzymes such as amylase and to meet present-day needs in the industrial sector. In the present study, a natural polymer gelatin is used with the nutrient agar medium to help in cell immobilization for maximum production of alkaline protease by strains of Bacillus. As alkaline protease has more applications than protease in industry, alkaline protease production from Bacillus is recommended. The highest amylase activity and proteolytic activity was given by B. subtilis coded by B3c and B. thuringiensis coded by B4i respectively. The highest yield organisms belonged to the nodules of leguminous plant.

## Data Availability

The data used to support the findings of this study are available from the corresponding author upon request.

## Authors’ Contributions

The budget for the experiment was incurred by the students as a part of their thesis work. Each participant designed the study and budget plan for all logistics, collected and analyzed the data, and participated in the draft and final write-up of the manuscript. Anup Basnet, the moderator and supervisor for this research participated in reviewing the initial and final drafts of the manuscript, data analysis, writing and interpretation of results, and final write-up of the manuscript. All authors reviewed and approved the final manuscript.

## Acknowledgments

We take this opportunity to express our profound gratitude and deep regards to our guide, lecturer and supervisor **Mr. Anup Basnet** for his guidance, monitoring, constructive suggestions and constant support to make this project a success. We appreciate our colleagues of St. Xavier’s College, Maitighar for discussions and communications for the formulation of the ideas and layout of this project. We also wish to express our gratitude to other staff members and lab technicians for their help and support. We are very much grateful to the principal **Father. Jiju Varghese, S.J.** and the head of department of Microbiology **Mr. Sudhakar Pant** for providing us the opportunity to embark on this project

We are grateful to the laboratory in-charge **Mr. Gopi Neupane** and **Ms. Anju Lama** for their kind co-operation during the research period.

## References

[1] R.M. Atlas and R Bartha. Microbial Ecology Fundamentals and Applications, 4th Edition. Pearson Education, pp 372–384, 2007.

[2] M.J. Pelczar, E.C.S. Chan and N.R. Krieg, Microbiology and Applied Based Approach, Third Reprint, Tata McGraw Hill Education Private Limited, pp 771–774, 2011.

[3] T.M. Madigan and J. M. Martinko, Brock Biology of Microorganisms, Pearson Education Limited, pp 374–594, 2006.

[4] A. Probanza, L. García, J. A. Palomino, M. R. Ramos and G. F. J. Mañero, “*Pinus pinea*, seedling growth and bacterial rhizosphere structure after inoculation with PGPR *Bacillus* (*B*. *licheniformis* CECT 5106 and *B. pumilus* CECT 5105)”, Applied Society and Ecology, 20:75–84, 2002.

[5] J. Harvey, K. P. Keenan and A. Gilmour, “Assessing biofilm formation by *Listeria monocytogenes* strains “, Food Microbiology 24: 380–392, 2007.

[6] C. B. Turnbull, Chapter 15, Bacillus Medical Microbiology, 4th edition, Baron S, editor. Galveston (TX): University of Texas Medical Branch at Galveston, 1996.

[7] S. K. Kochi, G. Schiavo, M. Mock and C. Montecucco, “Zinc content of the *Bacillus* anthracis lethal factor”, FEMS Microbiol Lett. 124:343, 1994.

[8] M. L. Prescott, P. J. Harley and A. D. Klein, Microbiology, 7th Edition, McGraw-Hill Limited, New York, USA, pp 964–976, 2008.

[9] N. A. Lyngwi and S. R. Joshi, “Economically important *Bacillus* and related genera: a mini review”, Biology of useful plants and microbes, 2014.

[10] G. Cercignani, M. C. Serra, C. Fini, P. Natalini, C.A. Palmerini, G. Mauni and P.L. Ipata, “Properties of a 5’-nucleotidase from *Bacillus cereus* obtained by washing intact cells with water”, Biochemistry, 13:3628–3634, 1974.

[11] D. A. Wood and H, Tristram, Journal of Bacteriology, 104:1045, 1970.

[12] P.V. Aiyer, “Amylases and their applications”, African Journal of Biotechnology, 4(8): 1525–1529, 2005.

[13] B. Pokhrel, P. Wanjaree and S. Singh, “Isolation, screening and characterization of promising amylase producing bacteria from sewage enriched soil.”, Int J Adv Biotechnol Res, 4 (2): 286–290, 2013.

[14] R. Gupta, Q. Beg and S. Khan Applied Microbiology and Biotechnology. 60: 381. 2003.

[15] L.S. Devi, P. Khaund and S.R. Joshi, “Thermostable α amylase from natural variants of *Bacillus* spp. prevalent in eastern Himalayan Range”, African Journal of Microbiology Research, 4(23):2534–2542, 2010.

[16] A. Edward, Nagula, I. D. McKelviea, P. Worsfold and S.D. Kolev, “The molybdenum blue reaction for the determination of 6 orthophosphate revisited: Opening the black box”, Analytica Chimica Acta. 890, 60–382, doi: 10.1016/j.aca.2015.07.030, 2015.

[17] K.S. Naidu,“Characterization and purification of protease enzyme”, Journal of applied pharmaceutical science, 01 (03):107–112, 2011.

[18] Monteiro de Souza P and P. Magalhães, “Application of amylase in industry – a review” Brazilian Journal of Microbiology, ISSN 1517-8382: 850–861, 2010.

[19] N. Akcan, “Production of extracellular alkaline protease from *Bacillus* subtilis RSKK96 with solid state fermentation, Eurasia J Biosci, 5:64–72, 2011.

[20] R. Banerjee and B.C. Bhattacharyya,“Extracellular alkaline protease of newly isolated *Rhizopus oryzae*”, Biotechnology letters, 14(4):301–304, 1992.

[21] P. Deb, A.M. Talukdar, K. Mohisna, P.K. Sarker and S.M.A. Sayem,“Production and partial characterization of extracellular amylase enzyme from *Bacillus amyloliquefaciens* P-001.”, Springer Plus, 2:154, 2013.

[22] T. P Keerthirathne, K. Ross, H. Fallowfield, and H. Whiley,“A Review of Temperature, pH, and Other Factors that Influence the Survival of *Salmonella*in Mayonnaise and Other Raw Egg Products”, Pathogens, 5(4), 63, 10.3390/pathogens5040063, 2016.

[23] S. Manandhar and S. Sharma, Practical Approach to Microbiology. 3rd Edition. National Book Centre. 2013.

[24] A. Pandey, P. Nigam, C.R. Soccol, V.T. Soccol, D. Singh and R. Mohan, “Advances in microbial amylases”, Biotechnol. Appl. Biochem, 31: 135–152, 2000.

[25] P. Bernfield. Amylases, - and In: Methods in Enzymology, Academic Press, New York, USA, 1: 149–158, 1955.

[26] T. Gibson and R. E. Gordon, Bacillus In Bergey’s Manual of Determinative Bacteriology. 8th Edition, Williams & Wilkins, Baltimore, pp 529–550, 1974.

[27] D. Daffonchio, N. Raddadi, M. Merabishvili, A. Cherif, L. Carmagnola, L. Brusetti and S. Borin,“Strategy for identification of *Bacillus cereus* and *Bacillus thuringienversis* strains closely related to *Bacillus anthracis*”, Applied and environmental microbiology, 72(2):1295–1301, 2006.

[28] I. Němečková, K. Solichová, P. Roubal, B. Uhrová and E. Šviráková, “Methods for detection of *Bacillus* sp., *B*. *cereus*, and *B*. *licheniformis* in raw milk”, Czech J. Food Sci. 29 (Special Issue): S55–S60 2011.

[29] F.A. Sharif and N.G Alaeddinoĝlu, “A rapid and simple method for staining of the crystal protein of *Bacillus thuringiensis**”,* Journal of Industrial Microbiology. 3: 227. 10.1007/BF01569580, 1988.

[30] D.K. Choudhary and B.N. Johri,“Interactions of *Bacillus* spp. and plants -with special reference to induced systemic resistance”, Elsevier, Microbiological Research. 64 (5): 493–513, 2008.

[31] V. Verma, M.S. Avasthi, A.R. Gupta, M. Singh and A. Kushwaha,“Amylase production and purification from bacteria isolated from a waste potato dumpsite in district Farrukhabad U.P State India’, European Journal of Experimental Biology, 1(3):107–113. 2011.

[32] K. Bose and D. Das,“Thermostable alpha-amylase production using *Bacillus licheniformis* NRRLB 14368”, Ind. J. Exp. Biol, 34: 1279–1282, 1996.

[33] Md.M. Hasan, L.W. Marzan, A. Hosna, A. Hakim and A.K. Azad,” Optimization of some fermentation conditions for the production of extracellular amylases by using Chryseobacterium and *Bacillus* isolates from organic kitchen wastes”, Journal of Genetic Engineering and Biotechnology, 15:59–68. 10.1016/j.jgeb.2017.02.009 2007.

[34] K. Suribabu, T. L. Govardhan and K.P.J. Hemalatha, “Influence of carbon sources on α-amylase production by *Brevibacillus* sp. under submerged fermentation”, International Journal of Research in Engineering and Technology, Volume: 03 Issue: 02, eISSN: 2319-1163 | pISSN: 2321–7308, 2014.

[35] S.K. Brar, M. Verma, R.D. Tyagi, R.Y. Surampalli, S Barnabé and Valéro JR, “*Bacillus thuringiensis* proteases: Production and role in growth, sporulation and synergism”, Process Biochemistry, Vol 4, PP 773–790, doi:10.1016/j.procbio.2007.01.015, 2007.

[36] G. Pant, A. Prakash, J.V.P. Pavani, S. Bera, G.V.N.S. Deviram, A. Kumar, M. Panchpuri and R.G. Prasuna, “Production, optimization and partial purification of protease from *Bacillus subtilis*”, Journal of Taibah University for Science, 9:50–55, 2015.

[37] D. Sarkar and G. Paul, “Extraction and Bio-chemical Characterization of Protease Enzyme from a Proteolytic bacteria Isolated from Dry Mixed Kitchen Waste”, International Journal of Current Microbiology and Applied Sciences, ISSN: 2319-7706, Volume 5 Number 3, PP 268–276, 2016.

[38] S. Sen and T. Satyanarayana, “Optimization of alkaline protease production by thermophilic *Bacillus licheniformis*. S-40”, Ind J Microbiol. 33:43–47, 1993.

[39] G.T Das and M.P Prasad, “Isolation, purification & mass production of protease enzyme from *Bacillus subtilis*”, International Research Journals of microbiology, 1(2):026–031, 2010.

[40] R.K Yadav, J.O. Baeg, G.H. Oh, N.J Park, K. Kong, J. Kim, D.W. Hwang and S.K. Biswas, “A Photocatalyst−Enzyme Coupled Artificial Photosynthesis System for Solar Energy in Production of Formic Acid from CO_2_”, J. Am. Chem. Soc. 134:11455–11461, 10.1021/ja3009902. 2012.

[41] Y. Li and A. A. Yousten, “Metalloprotease from *Bacillus thuringiensis*”, Appl. Microbiol, 30:352–361, 1976.

[42] M. R. Swain, S. Kar, G. Padmaja and R. C. Ray, “Partial Characterization and Optimization of Production of Extracellular”-amylase from *Bacillus subtilis* Isolated from Culturable Cow Dung Microflora”. Polish Journal of Microbiology, 55(4), 289–296,.2006.

[43] H.S. Alnahdi, “Isolation and screening of extracellular proteases produced by new isolated *Bacillus* sp.”, Journal of Applied Pharmaceutical Science, 2 (9):071–074, 2012.

[44] L. Szilard, Proc. U.S. Nat. Acad. Sci., 46, 277, 1960.

